# Personalized chordoma organoids for drug discovery studies

**DOI:** 10.1101/2021.05.27.446040

**Authors:** Ahmad Al Shihabi, Ardalan Davarifar, Huyen Thi Lam Nguyen, Nasrin Tavanaie, Scott D. Nelson, Jane Yanagawa, Noah Federman, Nicholas Bernthal, Francis Hornicek, Alice Soragni

## Abstract

Chordomas are rare tumors of notochordal origin, most commonly arising in the sacrum or skull base. Primary treatment of chordoma is surgery, however complete resection is not always feasible due to their anatomic location, and recurrence rates remain high. Chordomas are considered insensitive to conventional chemotherapy, and their rarity complicates running timely and adequately powered trials to identify effective regimens. Therefore, there is a need for discovery of novel therapeutic approaches. Drug discovery efforts in chordoma have been mostly limited to cell line models. Patient-derived organoids can accelerate drug discovery studies and predict patient responses to therapy. In this proof-of-concept study, we successfully established organoids from seven chordoma tumor samples obtained from five patients presenting with tumors in different sites and stages of disease. The organoids recapitulated features of the original parent tumors and inter-as well as intra-patient heterogeneity. High-throughput screenings performed on the organoids highlighted targeted agents such as PI3K/mTOR, EGFR, and JAK2/STAT3 inhibitors among the most effective molecules. Pathway analysis underscored how the NF-kB and IGF-1R pathways are sensitive to perturbations and potential targets to pursue for combination therapy of chordoma.

## Introduction

Chordoma is a rare malignant tumor that arises from the embryonic remnants of the notochord^1^. It typically affects older adults (median age 58.5), is more common in men than in women (5:3), and is diagnosed in about 300 Americans each year, with a median survival of just over 6 years^1,2^. There are three histological subtypes of chordoma: conventional, dedifferentiated and poorly differentiated^3–6^. Conventional chordoma accounts for the vast majority of cases; these are usually indolent, chemoresistant tumors^5,7,8^. The dedifferentiated subtype is reminiscent of high-grade pleomorphic spindle cell soft tissue sarcomas and typically follows an aggressive course^9^. Poorly differentiated chordoma is a rare, aggressive subtype affecting children and young adults and characterized by INI1 deletions^3,5^. Unlike conventional chordoma, dedifferentiated and poorly differentiated chordoma patients are typically treated with adjuvant chemotherapy^10^, with few documented responses^11,12^.

Treatment for chordoma relies primarily on surgery. Due to the anatomical location, complete resection can be challenging, particularly for clival tumors^8^. Even after achieving complete resection, recurrence rates remain high at approximately 40%^2^, often necessitating repeat surgeries. If the disease is metastatic or the patient is not a surgical candidate, there are few systemic treatment options available^10,13–15^. Traditional chemotherapeutic agents have not shown efficacy in this tumor type^7,8,14^, and there is no preferred regimen for the treatment of either locally recurrent or metastatic chordoma as of March 2021^10^. A small number of targeted agents have shown limited benefits in trials and are NCCN recommended for delaying tumor growth in some patients. These include imatinib with or without cisplatin or sirolimus, dasatinib, sunitinib, erlotinib, sorafenib and lapatinib for EGFR-positive chordoma^6^. A phase II trial of imatinib in 56 patients showed a 70% rate of stable disease at 6 months^15^. Sorafenib was associated with a progression-free survival (PFS) of 9-month in 73% of the 27 patients treated in a phase II trial^16^. The SARC009 study included 32 patients with unresectable chordoma treated with dasatinib, and showed a 54% PFS at 6 months^17^. However, most of these randomized trials have only extended PFS rather than achieving a partial or complete response^16^. Thus, there remains a significant need to identify efficacious therapies for chordoma^17^.

A substantial limitation that continues to hinder the identification of novel therapeutic avenues is the small number of validated chordoma models for pre-clinical research. Few immortalized chordoma cell lines have been reported^18–20^. While helpful, cell lines often fail at recapitulating the heterogeneity of the underlying disease and can deviate substantially from the parental tumor, resulting in changes to drug response^21^. As with most slow-growing tumors, the generation of patient-derived xenograft (PDX) models has lagged for chordoma, with moderate progress in recent years^20,22–26^. An approach to routinely establish chordoma organoids from biopsies or surgical specimen has the potential to power discovery studies to advance our understanding chordoma and identify new interventions^23,27^.

Patient-derived tumor organoids (PDOs) are ideally suited for modeling rare cancers and investigating their heterogeneity and drug response^27–33^. We have developed a high-throughput screening platform to test the response of patient-derived tumor organoids to hundreds of therapeutic agents, with results available within a week from surgery^28,34^. Here, we apply our approach and platform to generate and screen chordoma PDOs established from different tumor sites and histologies^28,34^. Determining the clinical efficacy of any new therapeutic approach is challenging in chordoma due to its natural history and slow growth rates. Personalized chordoma models can be rapidly established and effectively screened *ex vivo* to identify pathways sensitive to perturbations.

## Results

### Chordoma Patient Characteristics

In this proof-of-principle study, we aimed to determine whether clinically relevant, viable tumor organoid models can be established routinely from chordoma samples. We obtained n=7 samples from 5 patients enrolled in the study (**Table 1**). Patients were 60:40 male to female with a median age at diagnosis of 61, ranging from 27-to 73-year-old. Three patients were diagnosed with primary conventional chordomas (CHORD001, CHORD002, CHORD004), one with recurrent chordoma (CHORD003) and one with metastatic chordoma (CHORD002).

**Table 1.** List of patients and tumor sample characteristics. Collected attributes for each sample include the patient age at time of diagnosis, sex, tumor location, type of tissue procured, patient diagnosis at time of procurement, pathology report of samples, history of systemic treatment of patient, and the current disease status after follow up duration.

| Patient # | Age* | Sex | Tumor Location | Tissue Procured | Diagnosis** | Pathology | Systemic Treatment | Follow up |
| --- | --- | --- | --- | --- | --- | --- | --- | --- |
| CHORD001 | 64 | F | T3 vertebrae | Surgical resection of primary tumor | Primary conventional chordoma | 1.5 cm, differentiated with no necrosis | No | Disease free at 28 months |
| CHORD002 | 27 | M | Pelvis | Biopsy of pelvis metastasis (a) | Metastatic conventional chordoma, metastases in bone, liver and lungs | N/A | No | Progressive disease |
|  |  |  | L3 Vertebrae | Surgical resection of L3 tumor metastasis (b) | Metastatic conventional chordoma, metastases in bone, liver and lungs | 1 cm, 15% Ki-67, 15% necrosis | Yes | Progressive disease |
|  |  |  | Thoracic/Cervical Vertebrae | Resection of thoracic/cervical spine metastases (c) | Metastatic conventional chordoma, metastases in bone, liver and lungs | 35% Ki-67 | Yes | Progressive disease |
| CHORD003 | 61 | F | Sacrum | Surgical resection of recurrence | Recurrent chordoma, NOS | 3.4 cm, significant Ki-67 positivity | No | Residual disease after 4 months |
| CHORD004 | 48 | M | L1-L3 vertebrae | Surgical resection of primary tumor | Primary conventional chordoma | 6 cm, 20% necrosis | No | Disease free at 14 months |
| CHORD005 | 73 | M | Sacrum | Surgical resection of primary tumor | Primary chordoma, NOS | 11 cm, focal necrosis | No | Disease free at 12 months |

\* Age at time of diagnosis
\*\* Diagnosis at time of surgery

We obtained a single sample for each patient at the time of surgical resection, with the exception of CHORD0002, for which we procured n=3 samples: one from a biopsy (CHORD002a), a second from a surgical resection (CHORD002b) and the third from a subsequent spine metastasectomy (CHORD002c). Anatomically, out of the seven samples we procured, n=2 originated in the sacrum (CHORD003 and CHORD005), n=1 in the pelvis (CHORD002a), and n=4 from the vertebrae: T3 (CHORD001), L3 (CHORD002b), thoracic/cervical vertebrae (CHORD002c) and L1-L3 (CHORD004).

CHORD001 was diagnosed as a conventional type chordoma of the thoracic spine in a 64-year-old woman (**Table 1**). The primary mass was 1.5 cm in greatest dimension, moderately differentiated, with no necrosis or lymphovascular invasion. Histologic staining showed mixed staining for brachyury with a subset of the cells staining negative for this marker (**Figure 1**). The patient remains disease free at 28 months of follow up.

**Figure 1.**
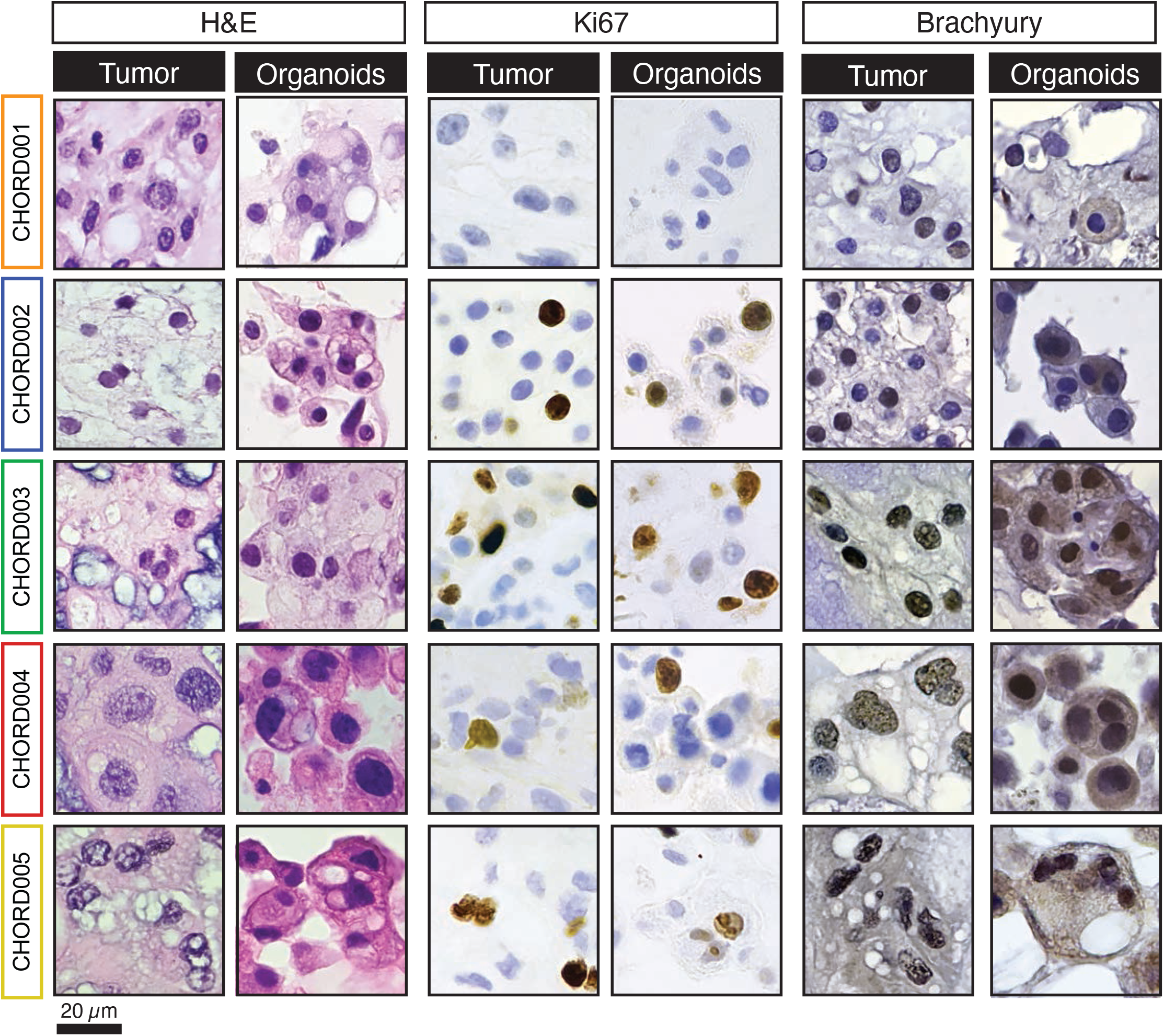
Histology and immunohistopathology characterization of chordoma samples and derived organoids. Formalin-fixed paraffin embedded sections from both the parent tumor and organoid were stained with H&E, Ki-67, and Brachyury. All organoids recapitulated features of the parent tumor. The sample shown for patient CHORD002 is CHORD002a. Scale bar 20 µm.

CHORD002 is a rare case of aggressive, metastatic conventional type chordoma of the sacrum diagnosed in a 27-year-old male (**Table 1**). The patient had a primary resection of an 8 cm mass with conventional histopathology and no lymphovascular invasion or necrosis. Next-generation sequencing analysis showed that the tumor had a low microsatellite instability (MSI), with loss of CDKN2A and CDKN2B. Only 6 months after surgical resection, the patient presented with metastatic recurrence to the lungs, spine, and pubic bone. A first sample was obtained from a biopsy of the pubic bone metastasis (CHORD002a) and confirmed to be conventional chordoma by pathologic morphology. The patient was treated with nivolumab, a PD-1 targeting monoclonal antibody^35^, in combination with the mTOR inhibitor nab-rapamycin^35^ as part of a clinical trial and started on a monoclonal antibody targeting RANKL, denosumab^35^. Treatment was discontinued after 3 cycles and followed by a combination regimen with trabectedin, an alkaloid that binds to DNA to arrest transcription^35^, nivolumab and an oncolytic herpes virus derivative^35^ as part of a different trial.

We then obtained a second sample from the resection of a vertebral metastasis (CHORD002b). The mass measured 1 cm in the largest dimension and was positive for INI1 and Ki-67. Most of the tumor was viable, with only 15% necrosis. The patient subsequently received one cycle of nivolumab, followed by a trial of DeltaRex-G, a retroviral vector encoding a cyclin G1 inhibitor^36^, followed by cetuximab, a monoclonal antibody targeting EGFR^35^ in combination with nivolumab in the setting of a clinical trial.

We procured a third tissue sample, CHORD002c, from a resection of metastatic lesions in the cervical and thoracic spine. This tumor was again classified as a conventional type chordoma with positive Ki67 (35%), retained/positive INI1, and positive brachyury. Whole exome NGS of the metastatic thoracic sample showed that loss of CDKN2A and CDKN2B was maintained, and the tumor had low mutational burden (TMB, 0.5 m/Mb). The patient subsequently received cyclophosphamide (an alkylating chemotherapeutic nitrogen mustard^35^), palbociclib (a small molecule inhibitor of CDK4/6^35^), and cisplatin (a platinum derivative that binds DNA^35^).

CHORD003 is a 61-year-old woman diagnosed with a not-otherwise-specified (NOS) chordoma of the sacrum (**Table 1**). At the time of resection, the mass was 8 cm in size, well-differentiated, with rare mitoses, and areas of focal necrosis. There was a first local recurrence 3 years post diagnosis in the right acetabulum. The mass was a low-grade conventional chordoma measuring 3 cm in size. We obtained a tissue sample from a second recurrence diagnosed 15 months later in the right acetabulum. The specimen measured 3.4 cm in size, from which we obtained a sample for our study (CHORD003, **Figure 1**). Immunohistochemistry showed retained INI1 and negative PDL-1 in tumor cells as well as significant Ki-67 positivity (**Figure 1**). MRI of the sacrum approximately after 4 months of follow up showed residual disease in the acetabulum.

CHORD004 is a case of NOS chordoma of the lumbar spine in a 48-year-old male (**Table 1**). We obtained a specimen from the initial resection the mass affecting the L1-L3 vertebrae (CHORD004). The tumor was 6 cm by radiographic imaging and mildly necrotic (20%). There is no evidence of disease at 14 months of follow up.

CHORD005 is a 73-year-old diagnosed with NOS chordoma of the sacrum (**Table 1**). The resected specimen measured 11 cm in the largest dimension, with negative margins and focal areas of necrosis. The patient has no evidence of recurrence after 12 months of follow up.

### Chordoma organoids establishment and characterization

We set to determine whether viable, tractable chordoma organoid models can be routinely established from tissue obtained from different surgical procedures (primary resection, metastasectomy, biopsy, **Table 1**), anatomical sites (cervical, thoracic or lumbar vertebrae, sacrum and pelvis) and disease characteristics (primary, recurrent or metastatic chordoma).

Tumor tissue was fast tracked to the lab for dissociation into a cell suspension containing single cells and small cell clusters (**Figure 2**). We obtained sufficient cells in all cases, including a biopsy (CHORD002a), and established viable organoids for characterization and screening for all n=7 samples (**Figure 1 and 2**). After seeding cells in a ring format according to our published protocols^28,34^ (see also **Materials and Methods**), we incubated them in serum free medium for a total of five days and imaged them daily (**Figure 2**)^28,34^. Individual PDOs showed a variety of behaviors in culture, with CHORD003 exhibiting robust growth, while CHORDO004 and CHORD005 were more indolent and mostly rearranged without sustained proliferation (**Supplementary Video 1**). CHORD002 had several vacuolated chordoma cells that assumed a spindle-like morphology and migrated in culture (**Figure 2**). To note, CHORD003 was obtained from a clinically rapidly growing recurrence, while CHORD002 was established from a diffusely metastatic chordoma, features that are reminiscent of the observed growth patterns.

**Figure 2.**
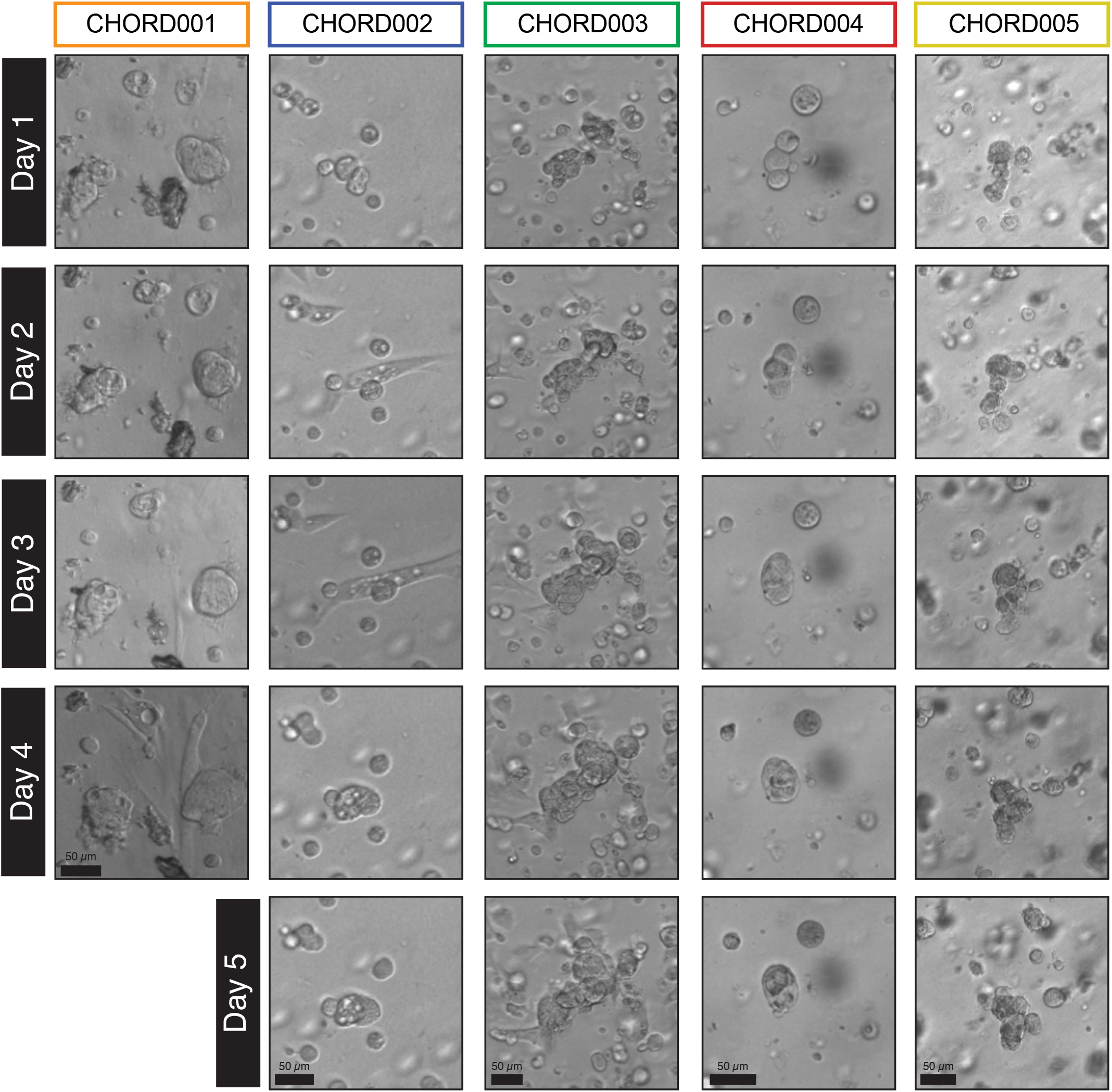
Morphology of the PDOs established as visualized by brightfield imaging of maxi-rings in 24-well plates. CHORD002-5 were imaged daily over a 5-day incubation period while CHORD001 was imaged for 4 days. The organoids displayed morphological features consistent with chordoma such as vacuolated cells arranged in clusters or nests. The sample shown for patient CHORD002 is CHORD002a. Scale bar 50 µm.

In order to better investigate features of the chordoma PDOs, we fixed and embedded cells for downstream analysis. When compared to the histopathology and immunohistochemical staining patterns of the parental tumors, PDOs retained all major features (**Figure 1**). For instance, both patient tumors and derived organoid cells show similar arrangements with cells in clusters and abundant eosinophilic vacuolated cytoplasm, as well as round nuclei containing small single or multiple nucleoli (**Figure 1**).

Matched chordoma parental tumors and organoids exhibited similar staining patters (**Figure 1**). We performed immunohistochemical staining for Ki-67, a marker of cell proliferation^37^. We observed positive Ki-67 nuclear staining in both tumor and organoid samples in all cases except for CHORD001. CHORD001 did not express Ki-67 in regions of the tumor as well as PDOs (**Figure 1**). Of note, CHORD003 had an overall higher percentage of Ki-67 positive cells in the tumor of origin, in line with the clinical characteristics of this tumor (**Table 1**). This feature is conserved in CHORD003 PDOs (**Figure 1**). A correlation between high Ki-67 positivity and aggressive behavior has been reported in chordoma^37^.

Next, we investigated expression of brachyury, a protein involved in notochordal development and a well-established chordoma marker^38^. CHORD003, CHORD004 and CHORD005 organoids and their parent tumor were positive for brachyury staining. On the other hand, CHORD001 was largely brachyury-negative and CHORD002 weakly positive for the parental tumor, and largely negative in the organoids (**Figure 1**). Overall, our tissue processing and culturing conditions yielded viable chordoma organoids with a 100% success rate (7/7) and immunohistopathology characteristics that mirror those of the patient tumor they are derived from.

### High-throughput drug screening of chordoma PDOs

Large scale screenings of chordomas have been few and limited to immortalized cell lines so far^39– 41^. To validate that our PDO mini-ring platform^28,34^ is suitable for high-throughput screening of chordomas, we performed proof-of-principle single concentration drug discovery screenings of up to 230 compounds on organoids established from CHORD001-5. Cells seeded in mini-ring format in 96 well plates^28,34^ were incubated for 3 days followed by drug treatments every 24 hours over two consecutive days. Viability was measured 24 hours after the last treatment by ATP release assay following our established protocols^28,34^.

We compared normalized viability and Z score results for the subset of overlapping drugs (n=124) tested on all seven samples (**Figure 3**). We used a distance matrix to perform hierarchical clustering of the tested drug on the basis of target similarity as annotated in the PubChem database^42^. Clinical samples were clustered by Euclidean distance of their drug response profiles (**Figure 3**). Dot maps showing additional molecules tested are visible in **Figure 4** and **Supplementary Figures 1-3**.

**Figure 3.**
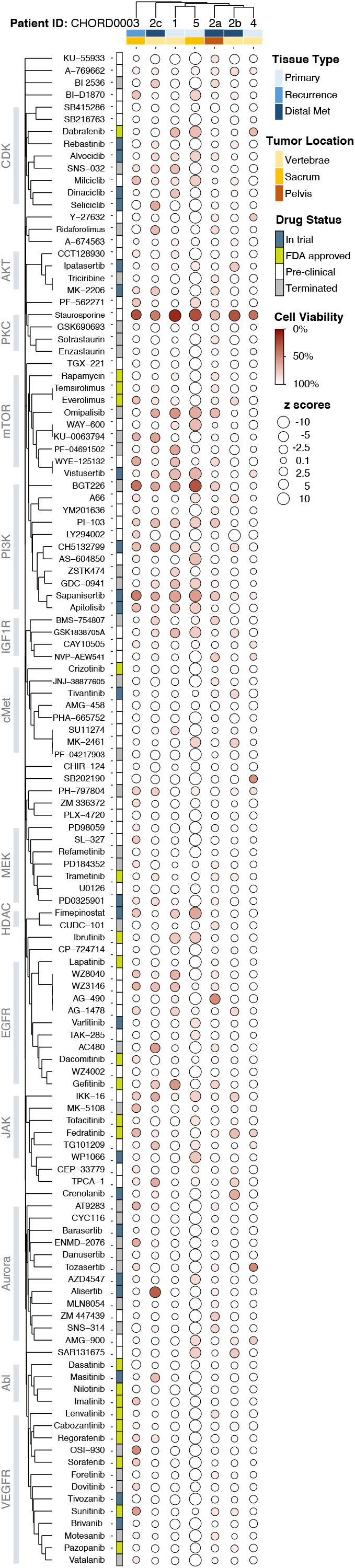
Dot map of high-throughput screening results for 124 overlapping small molecules tested on all PDOs. The size of each circle represents the Z score, with larger ones indicating a higher Z score value. The color of each point represents the normalized cell viability %. The drugs are clustered using the Jaccard distance based on common protein targets, and cases are clustered based on their similarities in drug responses. Covariates included represent the type of tissue and location of each sample as well as FDA status for the various drugs.

**Figure 4.**
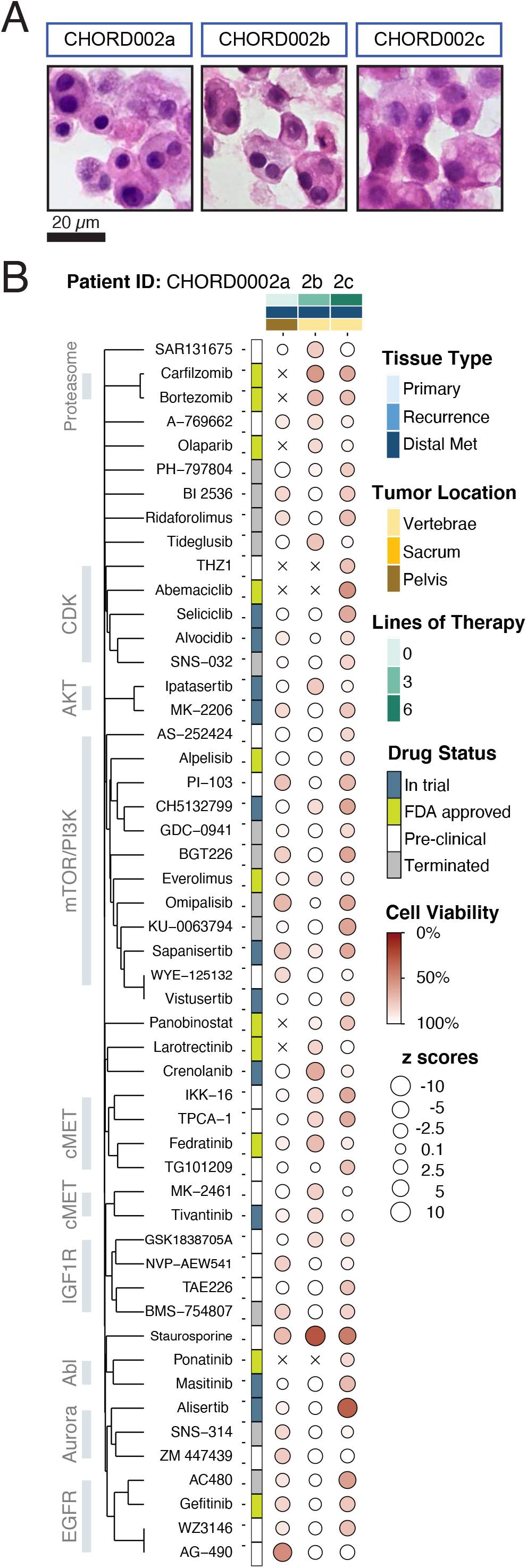
Dot map of drug screening on all three samples procured from patient CHORD002. Drugs that induce 15% cell death in at least two of the samples are visualized (51 total). The size of each circle represents the Z score, with larger ones indicating a higher Z score value. The color of each point represents the normalized cell viability %. The drugs are clustered using the Jaccard distance based on common protein targets. Samples are ordered by date of acquisition. Covariates included represent the type of tissue, anatomic location of each sample and the number of lines of treatment the patient was exposed to prior to procurement of sample as well as FDA status at the time of publication.

Overall, the chordomas we tested showed marginal level of responses, with only the positive control staurosporine, a potent multi-kinase inhibitor, showing consistent albeit moderate efficacy (9% - 71% residual cell viability) across all samples (**Supplementary Table 1**). For five out of n=7 samples tested, our positive control (staurosporine) was the only drug with a measured residual cell viability ≤ 25%. Response rates defined as residual cell viability ≤ 25% varied between 0.4 - 3% (median 0.65%) including staurosporine and 0% - 2.4% excluding the control. This is markedly lower than the responses we observed for ovarian cancer PDOs we tested following the same protocol (0.8% - 6%, median 1.9%, n=4, excludes staurosporine)^28^, and in line with clinical findings of high therapeutic resistance of chordomas^43^.

Given the generally low response rates, we investigated drugs causing residual cell viability of ≤ 50% and ≤ 75% (**Supplementary Table 1**). Response rates were 0% - 6% for the first group of drugs inducing at least 50% cell death (median 1%) and 2% 29% for the second, causing at least 25% cell death (median 8%, **Supplementary Table 1**). We observed sensitivity toward inhibitors targeting the PI3K and mTOR pathways, specifically for samples CHORD001, CHORD005, CHORD003 and CHORD002c with an average residual cell viability between 60% (CHORD001) and 71% (CHORD002c). Molecules such as BGT226, sapanisertib, omipalisib and vistusertib were effective in inducing ≥ 25% cell death on average (**Figure 3, Supplementary Table 2**). Validation screenings for sapanisertib and several other molecules tested at concentrations between 0-10 µM largely confirmed the results of our discovery screenings (**Supplementary Figure 1A-B**).

Beside mTOR/PI3K inhibitors, the JAK2-targeting molecule fedratinib also showed partial efficacy inducing ≥ 20% cell death in 3/7 samples (CHORD002b, CHORD003, CHORD004, **Supplementary Table 2**). Inhibition of the JAK2/STAT3 pathway in vitro has been shown to reduce viability of chordoma cell lines^44,45^. Gefitinib, an FDA approved EGFR inhibitor, showed responses in 3/7 samples (CHORD002a and c, CHORD005, **Supplementary Table 2**). Gefitinib has been highlighted as a potential effective agent against chordoma on large-scale screenings of chordoma cell lines^40^.

The Abl-targeting drug imatinib and multi-tyrosine kinase inhibitor sunitib showed efficacy in a single sample, CHORD003, with 27% cell death, Z score -1.6 and 44% cell death, Z score -2.7 (**Supplementary Table 2**). Both molecules are NCCN recommended as systemic therapy for some chordomas^13^.

### Drug response evolution of CHORD002

Metachronous procurements of tumors from one patient with metastatic chordoma, CHORD002, allowed us to investigate the evolution of drug responses over time and therapy pressure (**Figure 4** and **Supplementary Figure 2**). All organoid samples had similar histology (**Figure 4a**). CHORD002a organoids were established from a biopsy of a pelvic metastatic lesion obtained prior to the initiation of any systemic therapy. This sample displays moderate sensitivity to drugs of the mTOR family and EGFR families (**Figure 4**).

CHORD002b organoids were generated from tissue obtained at the time of surgical resection of a lumbar spine metastasis. This specimen was obtained after 3 lines of systemic treatment that included immune checkpoint blockade and the mTOR inhibitor rapamycin, as well as immune checkpoint blockade in combination with trabectedin^35^, and palliative radiotherapy to the lumbar spine. The original tumor sample of CHORD002b showed 15% necrosis on pathology examination and the organoid established from this sample showed diminished sensitivity to the mTOR family of drugs (**Figure 4**).

Lastly, we established CHORD002c organoids from a surgical resection of cervical/thoracic spine metastasis, obtained after an additional 3 lines of treatment with EGFR-targeting monoclonal antibody cetuximab in addition to immune checkpoint blockade, and a trial of the retroviral vector DeltaRex-G. Interestingly, the organoid established from this sample did not show increased sensitivity to AC480, gefitinib and WZ3146. In addition, sample CHORD002c showed restored sensitivity to the mTOR-targeting drugs (**Figure 4**).

### Biological pathways associated with metabolism, bone signaling and inflammation are sensitive to perturbation in chordoma PDOs

The data collected over a large chemical space allowed us to evaluate which pathways were most affected in our screenings. To do this, we compiled a comprehensive list of known targets for each drug we screened according to the Pubchem^42^ database and overlayed these with well-characterized pathways described by WikiPathways^46^.

For each sample, we generated a matrix of drugs and corresponding protein targets weighted by the measured cell viability (see **Materials and Methods**). Using WikiPathways^46^, we compiled a pathways-protein matrix, which was overlayed with the drug-target interaction matrix to investigate the effect of the tested drugs on these pathways and generate a pathway score. We then selected the top 30 scored pathways in each sample, and visualized those shared between a minimum of two chordomas in **Figure 5**, with the size of each circle represents the pathway score, and the color representing the proportion of the genes in a pathway that are targeted by the tested drugs. The pathways are clustered based on protein overlap (**Figure 5**).

**Figure 5.**
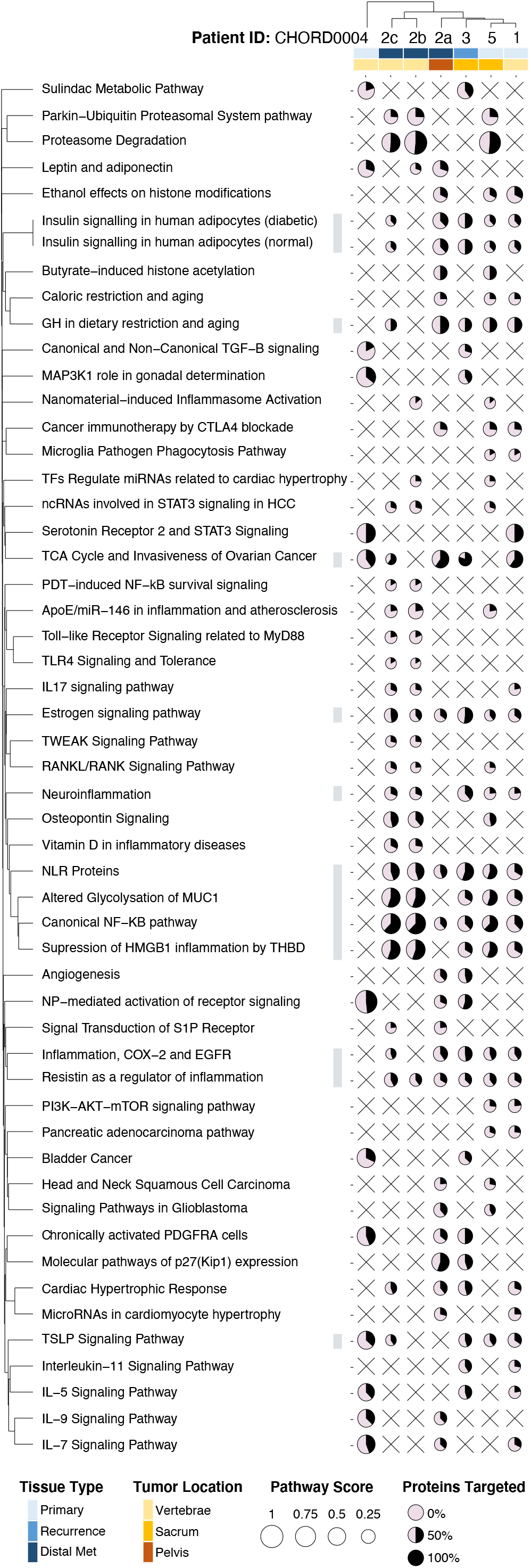
Pathway analysis for all chordoma PDOs. The top 30 ranked pathways that are affected in at least 2 samples are visualized (51). The color of each dot corresponds to the percentage of proteins that were targeted while the size corresponds to the pathway score. The pathways are clustered using the distance based on their shared proteins. The samples are clustered by the Euclidian distance of their pathway scores. Covariates included represent the type of tissue, and site of each sample.

Our analysis highlighted how metabolism-related pathways were mostly sensitive to perturbation, with drugs targeting these pathways reducing chordoma PDOs viability in most samples. Affected pathways include insulin signaling (5/7 samples), TCA cycle (5/7 samples), and estrogen signaling pathways (6/7 samples) (**Figure 5**). Interestingly, bone signaling such as the RANK/RANKL and osteopontin signaling pathways were also impacted in 3/7 samples (**Figure 5**).

Additionally, pathways related to inflammation, and involving NF-kB, MUC1, HMGB1 and resistin were affected in most of our samples (> 5/7, **Figure 5**). Targeting the proteasome pathway had also a profound effect in 3/7 samples; we believe this is due to the large effect of the drugs bortezomib and carfilzomib that were screened on samples CHORD002b, CHORD002c, and CHORD005, and tend to elicit large in vitro responses (**Figure 5**).

## Discussion

Chordoma remains an understudied rare cancer with limited therapeutic options and few patient-specific models^47^. We propose an approach to develop personalized chordoma organoid models for biological characterization and high-throughput drug screenings from tissue obtained through a variety of surgical procedures and tumor sites. The chordoma organoids we have grown recapitulate features of the original tissue, such as histopathology, positivity for brachyury and rate of Ki-67 staining (**Figure 1**), as well as demonstrate growth patterns that are representative of specific clinical features (**Figure 2**). While the PDO models shared the immunohistopathological profile of their parent tumor sample, they maintained individual differences between patients. For instance, CHORD002 is a very aggressive metastatic chordoma with widely disseminated metastases and cells had unique migration patterns in culture (**Figure 2, Supplementary Video 1**). Furthermore, patient CHORD003 developed multiple local recurrences. The organoids established from CHORD003, showed robust proliferation by growth pattern (**Figure 2**) and Ki-67 staining (**Figure 1**). These observations suggest that our models may be integrated in the framework of personalized medicine.

The PDOs we developed through our established platform^28,34^ can be effectively screened to identify promising drugs and sensitive pathways (**Figure 3 - 5**). Drugs targeting some of the pathways we identified have been evaluated in trials. For instance, PI3K/ mTOR inhibitors have been evaluated in chordoma both as monotherapy as well as in combination with imatinib^48^. The PI3K/mTOR targeting drugs sapanisertib and vistusertib showed efficacy in our organoid screenings (**Figure 3**). Both drugs have been found to be effective in chordoma cell lines^23,49^. Our screening additionally identified other PI3K/mTOR targeting drugs such as apitolisib and omipalisib. The clinical efficacy of these drugs in chordoma remains to be determined in trials.

EGFR activation, measured by phosphorylation, is frequently seen in chordomas^50^. EGFR targeting drugs such as lapatinib have shown some efficacy in controlling disease and are approved for use in EGFR positive chordomas^10,51^. Additional EGFR targeting drugs that have been tested in clinical trials for chordoma include erlotinib, linsitinib, afatinib, and the monoclonal antibody cetuximab^48,49^. We observed moderate efficacy of the EGFR targeting drug gefitinib in 3/7 samples. While gefitinib has not been tested in clinical trials as monotherapy in chordoma, it has been tested in combination therapy with cetuximab. In 2 patients the combination of cetuximab and gefitinib showed a partial response with a duration of 9 months for a single patient^52^ and a reduction of tumor bulk by 44% in another^53^. In vitro screenings on chordoma cell lines have also identified gefitinib as an effective EGFR targeting drug^40^.

The STAT3 transcriptional pathway has been found to be implicated in the development of chordomas^44,54^. The efficacy of JAK2/STAT3 targeting drugs on chordoma has only been evaluated in in vitro studies that showed efficacy on chordoma cell lines^45,55^. We found the JAK2 targeting drug fedratinib to be effective on a subset of our samples (3/7, **Figure 3**). Fedratinib has not been tested clinically in chordoma yet.

Our analysis identified pathways that are likely to be dysregulated in chordoma, such as the NF-kB pathway. For instance, IKK-16 (an inhibitor of NF-kB) was active in most of our samples (**Figure 3 and 5**). Other drugs that target the NF-kB pathway, such as IMD-0354, have also shown efficacy in chordoma xenografts^24^. We identified other pathways likely to be dysregulated and targetable in chordoma, such as the insulin signaling pathway (**Figure 5**). Previous studies have found increased expression of IGF-1R in chordomas^40^. Clinical evidence that this pathway can be targeted in chordoma includes a case report of a chordoma patient treated with the combination of the IGF-1R inhibiting drug linsitinib and the EGFR inhibitor erlotinib for 18 months, who achieved a partial response^56^. Further clinical evidence was seen in a phase 1 clinical trial of the same combinatorial regimen, with one chordoma patient experiencing sustained partial response^57^.

Our analysis also highlighted the MUC1 pathway, that promotes therapy resistance^58,59^ (**Figure 5**). Expression of MUC1 plays a protective role in tumors against immune attacks, and inhibition of MUC1 can overcome the resistance to immune processes^59^. The association between MUC1 and chordoma is believed to be mediated by brachyury, as MUC1 was found to be upregulated in tumors that highly express brachyury^59^. Overall, the identification of specific pathways highlights avenues for future combinatorial studies.

Finally, our observations from PDOs collectively confirm previous findings in chordoma patients and cell lines. Yet, it is noteworthy that PDOs were highly patient-specific, with characteristics differing not just between patients but also between samples obtained from the same patient at different time points and sites (**Figure 1-5**). Furthermore, none of the therapies identified worked on each and every one of the PDO models tested (**Figure 3**). This suggests that a personalized approach to the treatment of chordoma would allow better stratification of patients to efficiently target pathways that are susceptible to therapy in each case. In conclusion, our organoid-based functional sensitivity profile may be used to tailor therapy to each individual, both improving the efficacy of treatment as well as sparing patients ineffective therapies and their associated toxicities.

## Acknowledgments

We thank Dr. Robert Damoiseaux and the UCLA MSSR Core for providing part of drug libraries included in this study. This project was supported by the NIH R01CA244729 grant (to A.S.) and a Seed Grant from the UCLA DGSOM (to A.S., N.F. and J.Y.).

## Materials and Methods

### Tissue Procurement and Processing

Patients were consented on the UCLA IRB approved protocol 18-000980. Samples obtained in the operating room, were stored in RPMI and immediately transferred to the laboratory. Tumors were cut into small, 1-3 mm^3^ fragments and dissociated to single or small cell clusters by adding Collagenase IV (200 U/ml) and incubating at 37 °C with 5% CO_2_. Samples were vortexed every 15 minutes and cells collected every 2 hours. After red blood cells lysis, tumor cells were filtered through a 70 μm cell strainer and then counted^34^.

### High-throughput Drug Screening: Mini-rings establishment

We followed our established methods to generate organoids amenable to high throughput drug screening^28,34^. Briefly, a suspension of 5000 cells/well (single cells or small clusters) was plated at the rim of white 96-well white plates (Corning #3610) in a 3:4 mixture of media (StemCell Technologies # 05620) and Matrigel (BD Bioscience CB-40324). Plates were incubated at 37 °C with 5% CO_2_ for 30 min to solidify the gel before adding 100 µl of pre-warmed Mammocult medium to each well using a liquid handler. Three days after seeding mini-rings, cells were treated by replacing the media with fresh Mammocult containing the indicated drugs. The same procedure is repeated after 24 hours.

### High-throughput Drug Screening: ATP assay

Twenty-four hours after the last treatment, the media was removed and wells washed with 100 µl of pre-warmed PBS. Organoids were released from Matrigel by incubating at 37°C for 25 min in 50 µl of 5 mg/mL dispase (Life Technologies 17105-041). Plates were vigorously shaken at 80 rpm for 5 min prior to adding 75 µl of CellTiter-Glo 3D Reagent (Promega #G968B) to each well. Luminescence was measured after a total 30 minute incubation at room temperature using a SpectraMax iD3 (Molecular Devices) over 500 ms of integration time. For each 96 well-plate, 8 wells are dedicated to staurosporine positive controls and 8 wells for DMSO negative controls. The rest of the wells were used to plate the samples with varying drugs. The drugs are tested at single concentration (1 µM) for discovery screening as established in Phan et al, 2019^28^ or at 0.1, 1, and 10 µM in duplicates for validation screenings.

### Maxi-rings Preparation

Suspensions of single cells and small clusters were plated at the rims of 24 well plates (Corning #3527) in the same 3:4 mixture of media (StemCell Technologies #05620) and Matrigel (BD Bioscience CB-40324). For this set up, we plate 100’000 cells/well in a 70 µl mixture/ring as previously reported^28,34,60^. Plates were incubated at 37 °C with 5% CO_2_ for 30 min to solidify the gel before adding 1 ml of pre-warmed Mammocult medium to each well. Medium is fully removed and replaced with fresh pre-warmed one after an initial 3 day incubation period. After 5 days of growing, maxi-rings were washed with 1 ml pre-warmed PBS, and fixed in 500 μl of 10% buffered formalin (VWR #89370-094). Organoids were then transferred to a 15 ml falcon tube the following day, washed twice in PBS followed by the addition of 5 μl of Histogel (Thermo Scientific #HG-40000-012) and transfer to a cassette. We then proceed with standard embedding, sectioning and H&E staining.

### Immunohistochemistry

For Ki-67 staining, slides were baked at 45 °C for 20 min and deparaffinized in xylene followed by washes in ethanol and deionized water. Endogenous per-oxidases were blocked with Peroxidazed-1 (Biocare Medical #PX968M) at RT for 5 min. Antigen retrieval was performed in a NxGEN Deloaking Chamber (Bio-care Medical) using Diva Decloacker (Biocare Medical #DV2004LX) at 110 °C for 15 min for Ki-67/Caspase-3 staining. The combo Ki-67/Caspase-3 (Biocare Medical #PPM240DSAA) solution is pre-diluted and was added to the sample for 60 min at RT. Secondary antibody staining was performed with MACH 2 double Stain 2 (Biocare Medical #MRCT525G). All secondary antibodies were incubated at RT for 30 min. Chromogen development was performed with Betazoid DAB kit (Biocare Medical #BDB2004). The reaction was quenched by dipping the slides in deionized water. Hematoxylin-1 (Thermo Scientific #7221) was used for counterstaining. The slides were mounted with Permount (Fisher Scientific #SP15-100). For brachyury staining, an anti-brachury antibody (Abcam catalog number ab209665) was used at a 1/6’000 dilution and incubated for 50 minutes. The BOND-III Fully Automated IHC staining was detected using the BOND Polymer Refine Detection (Leica, DS9800). Images were acquired with a Revolve Upright and Inverted Microscope System (Echo Laboratories).

### Cell Viability Calculation

The luminescence values are normalized to the control DMSO vehicle wells, and Z scores are calculated as (viability of drug X – viability of vehicle) / SD of vehicle. For the dot map visualization and pathway analysis, we used drug viability values at 1 µM. PubChem and the FDA orange book databases were used to determine the FDA approval status of the screened drugs, results were categorized as either pre-clinical, in trial, FDA approved, or terminated in **Figures 3, 4** and **Supplementary Figures 2 and 3**.

### Target and Pathway Analysis

The BioAssay results from the PubChem database for each of the screened drugs were obtained using the PubChem API. To create a list of the most active targets for each drug, we selected only protein targets that are within 10-fold of the second lowest reported value for K_d_ or IC_50_ in the PubChem database. For drugs that did not have any BioAssay data in PubChem, protein targets were manually curated from the literature.

To perform the pathway analysis, we first generated a matrix of drug and protein target interactions *n*_drugs_ x *m*_protein_ with values of 0 or 1; with 1 indicating the presence of an interaction between a drug and a certain protein target, and 0 indicating its absence. We multiplied each row by a weight proportional to the mean viability of the organoids treated by each drug (1-(mean viability/100)). We then multiplied this matrix by a vector of 1’s to obtain a row-wise summation of the protein target-viability values. Subsequently, we normalized the row vector by dividing each element by the sum of non-zero column entries in the *n*_drugs_ x *m*_protein_ matrix. For example, if the total number of drugs that targets a given protein is 5, we divide the elements of the matrix corresponding to that protein by 5. This analysis produced a row-wise vector of weighed targets causing organoids viability changes.

We then mapped this list of proteins to the canonical pathways defined by the Wiki Pathway Database (version 20210210^46^). We used a subset of pathways that excluded those not biologically pertinent to cancer, such pathways related to microorganisms and pathogens, as well as the newly added COVID-related pathways. To create this map, we populated a new *n*_pathway_ x *m*_protein_ matrix with 1’s indicating the presence of a protein in a particular pathway or 0’s indicating its absence. We then normalized the rows in the *n*_pathway_ x *m*_protein_ matrix to account for the differences in the number of proteins included in each pathway. To obtain the relative effect that targeting a specific pathway has on the viability of each chordoma organoid case, we multiplied the *n*_pathway_ x *m*_protein_ mapping matrix by the normalized *n*_drugs_ x *m*_protein_ vector. The resulting vector represents the relative impact that targeting a given pathway has on the viability of the organoids.

### Dot Map Generation

We used R and the tidyverse package to plot the viability data or pathway data for each case and the arrays of drugs tested. We clustered the drugs (**Figure 3-4**) and pathways (**Figure 5**) by the number of overlapping elements. This was done using the Jaccard distance function in R. Cases that had similar responses to a given drug or similar affected pathways were clustered together using the distance function, as parameterized by a continuous Euclidean distance. The visualizations for all dot maps were produced using the ggplot2 package and finalized with Adobe Illustrator.

## Supplementary Material

### Supplementary Video

**Supplementary Video 1.**
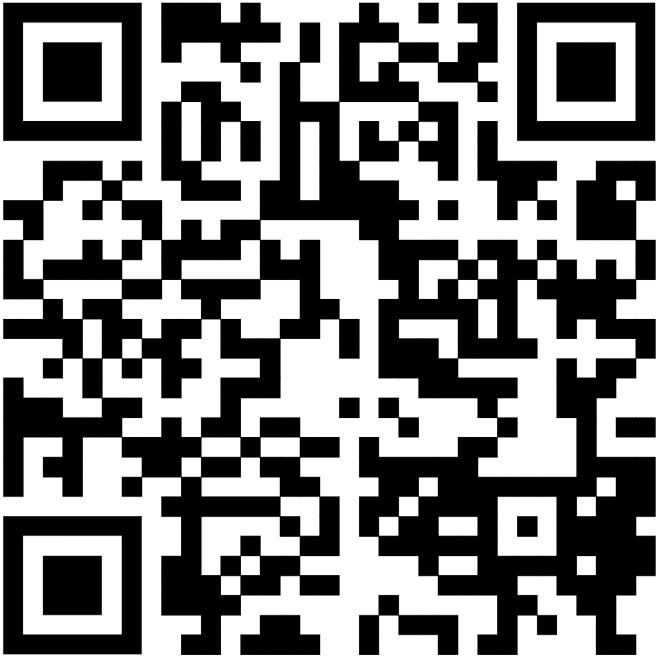
Compiled images as shown in Figure 2. Scan the QR code to access the video or visualize at the following link: https://youtu.be/5aOuyUMkpaE

### Supplementary Tables

**Supplementary Table 1.** Summary of results of the high-throughput screening for all PDOs. For each sample, the number of total drugs that were tested, and the percentage of drugs inducing 25%, 50% and 75% are listed.

| Sample # | Number of Drugs Tested | % Drugs inducing 25% Cell Death* | % Drugs inducing 50% Cell Death* | % Drugs inducing 75% Cell Death* |
| --- | --- | --- | --- | --- |
| CHORD001 | 140 | 7% (10/140) | 1% (1/140) | 0% (0/140) |
| CHORD002a | 154 | 2% (3/154) | 0% (0/154) | 0% (0/154) |
| CHORD002b | 165 | 8% (14/165) | 1% (2/165) | 0% (0/165) |
| CHORD002c | 188 | 24% (46/188) | 6% (11/188) | 3% (5/188) |
| CHORD003 | 231 | 15% (35/231) | 1% (3/231) | 0% (0/231) |
| CHORD004 | 228 | 7% (15/228) | 1% (2/228) | 0% (0/228) |
| CHORD005 | 171 | 29% (33/171) | 5% (8/171) | 0% (1/171) |

\*Excludes Staurosporine

**Supplementary Table 2.**
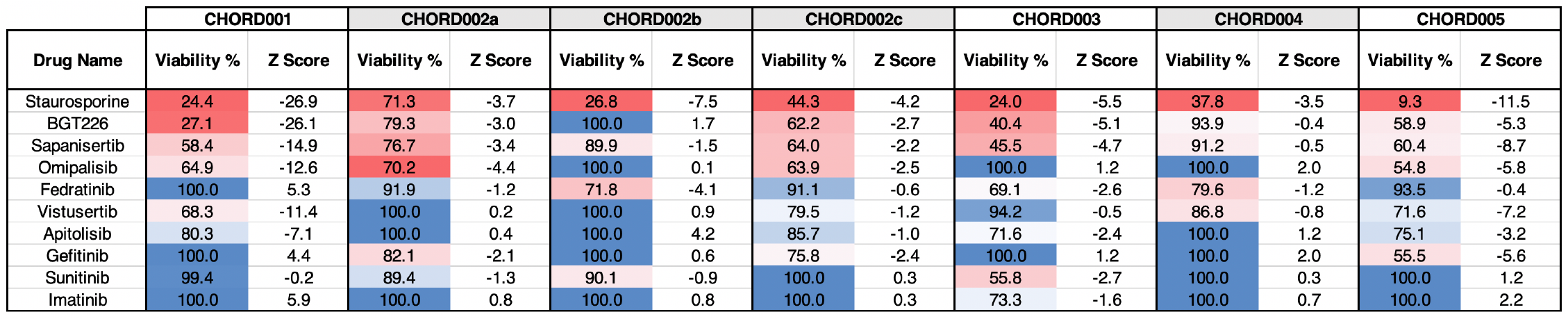
Summary of results of the high-throughput screening on a selected number of drugs. Viability and Z score are listed across all samples

### Supplementary Figures

**Supplementary Figure 1.**
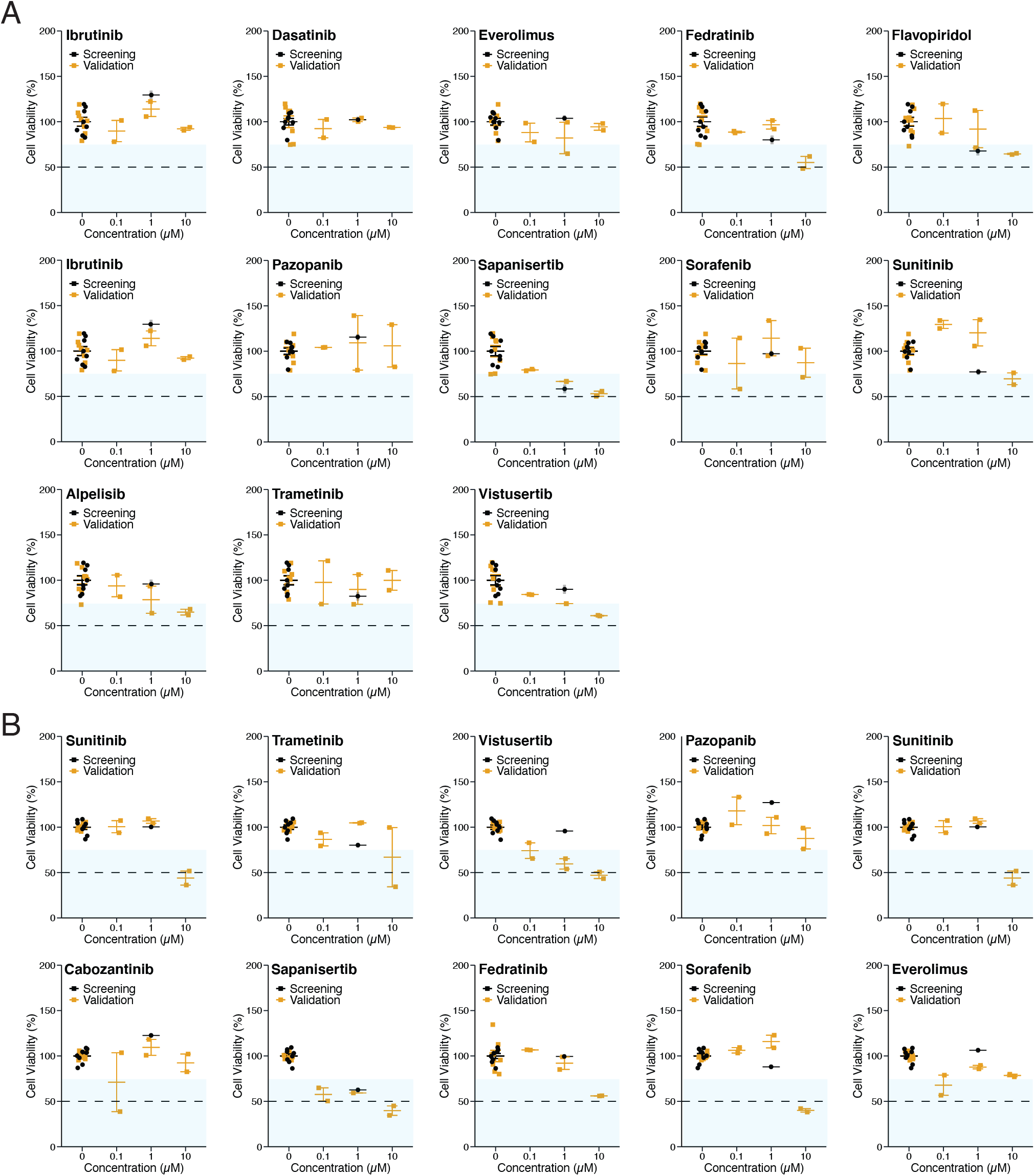
Dose-response plots of drugs tested in both single concentration (1µM) discovery screening and in subsequent validation screening at multiple concentrations (0.1µM, 1µM, and 10µM) on samples CHORD002c **(A)** and CHORD005 **(B)**. Points represent individual wells, error bars show SEM. Discovery screening data is shown in black while validation screening values are in yellow.

**Supplementary Figure 2.**
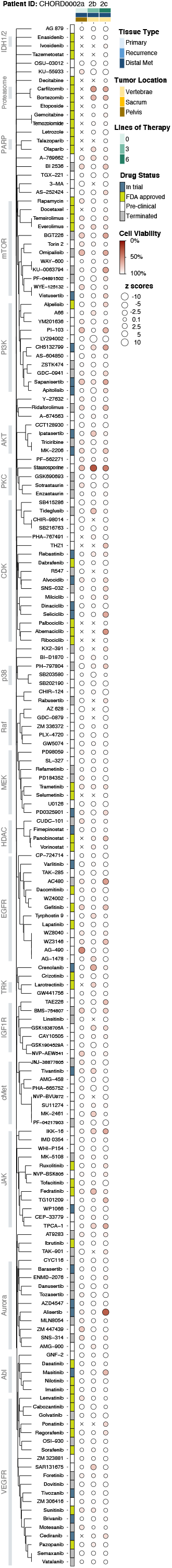
Dot map showing the full drug screening results on all three samples obtained from patient CHORD002. The number of drugs tested on sample CHORD002a, CHORD002b, and CHORD002c is 155, 159 and 179 respectively, with X labeling drugs that were not screened. The size of each circle represents the Z score, with larger ones indicating a higher Z score value. The color of each point represents the normalized cell viability %. The drugs are clustered using the Jaccard distance based on common protein targets. Samples are ordered by date of acquisition. The covariates included show the type of tissue, anatomic site and the number of lines of treatment the patient was exposed to prior to procurement of sample.

**Supplementary Figure 3.**
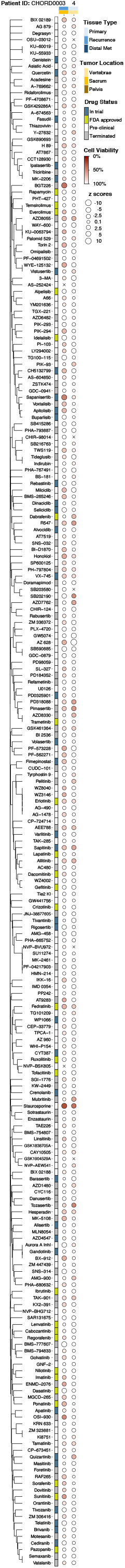
Dot map of the complete drug screening results on samples CHORD003 and CHORD004. Number of drugs tested on sample CHORD003, and CHORD004 is 232 and 229 respectively, with X marking compounds not screened in one of the samples. The size of each circle represents the Z score, with larger ones indicating a higher Z score value. The color of each point represents the normalized cell viability %. The drugs are clustered using the Jaccard distance based on common protein targets. Co-variates included represent the type of tissue, and anatomic location of each sample.

